# Assessment of polymeric immunoglobulin A and M via detection of the joining chain

**DOI:** 10.64898/2026.07.14.737820

**Authors:** Nienke Oskam, Jim Keijser, Marij Streutker, Sofie Keijzer, Pleuni Ooijevaar-de Heer, Gerard van Mierlo, Ninotska Derksen, T2B! immunity against SARS-CoV-2 study group, Gestur Vidarsson, Theo Rispens

## Abstract

Polymeric immunoglobulin M (IgM) and A (IgA) play key roles in systemic and mucosal immunity, yet quantitative assessment of their polymeric forms has been hampered by the lack of robust, high-throughput assays. Polymerization of both isotypes implies incorporation of the joining chain (J chain), making direct detection of integrated J chain an attractive surrogate marker. Here, we report the generation and characterization of a novel panel of monoclonal antibodies targeting human J chain. Binding analyses revealed distinct antibody clusters with differential preferences for IgA-J and/or IgM-J. We developed sensitive ELISAs that allow reliable quantification of J-chain– containing IgM and IgA in recombinant preparations and complex biological samples such as serum and saliva. For IgM, assay performance in serum required mild dissociation of the IgM– CD5L complex, enabling accurate detection of integrated J chain. For IgA, clone 9G10 showed remarkable specificity for IgA-J, with minimal cross-reactivity to IgM. Application of these assays demonstrates that on average, 10% of circulating IgA is J-chain–containing, with proportional contributions of IgA1 and IgA2, and enables high-throughput measurement of antigen-specific polymeric IgA responses, exemplified by SARS-CoV-2 vaccination. These tools provide a long-needed platform to study polymeric antibody dynamics in health, infection, vaccination, and B-cell–driven diseases.

## Introduction

Human antibodies are made up of five isotypes, of which both immunoglobulin M (IgM) and A (IgA) can be secreted as polymers (**Figure 1A**). This is because both isotypes contain a C-terminal tailpiece with unpaired cysteine residues that steer both the association with other monomeric subunits and the joining chain (J chain).^1,2^ The J chain is highly evolutionary conserved ^46^ with a unique tertiary structure that integrates into an IgM or IgA polymeric structure, forming two disulfide bonds with the unpaired cysteines of two different tailpieces.^47,48^

**Figure 1.**
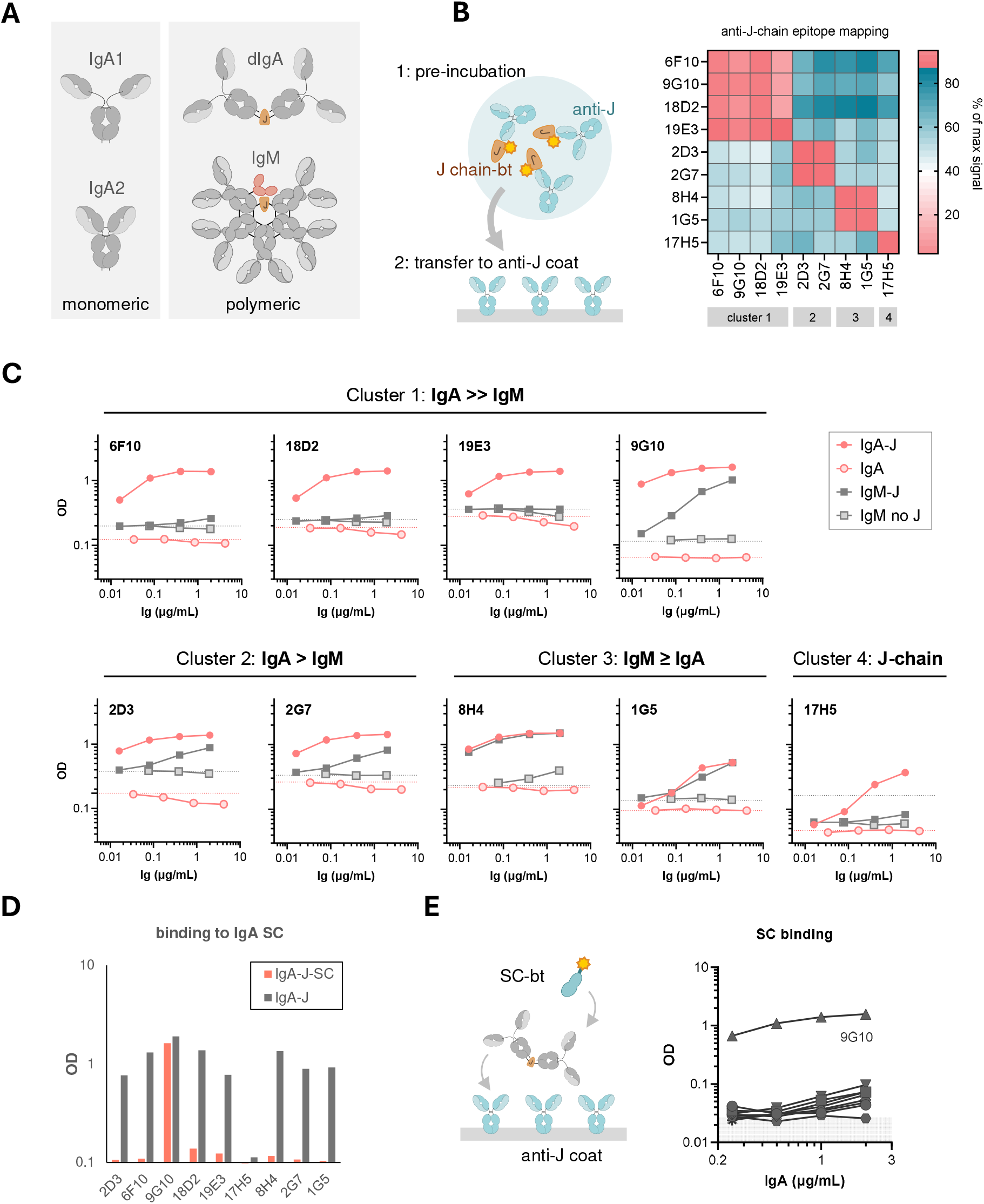
Global characterization of anti-J chain clones. A) Schematic of the structure of IgA and IgM-CD5L in circulation. Besides monomeric and dimeric IgA, larger polymers have also been described, which also contain a J chain. B) Overview of epitope mapping of anti-J chain clones. Biotinylated J chain was pre-incubated with one of the anti-J chain mAbs and then added to plates coated with one of the other anti-J chain mAbs, after which bound J chain was detected. No signal indicated that the pre-incubated clone and coated clone bind in a similar area on the J chain and thus means that they likely bind (overlapping) epitopes. C) Binding of anti-J chain clones to recombinant IgA1 (anti-spike clone 1-18) and IgM (ACPA clone 2D5) with and without J chain. Representative plots of 2-3 experiments. D) The ability of the clones to bind to secretory antibodies was tested. Capture of 0.25 ug/mL of IgA1-J or IgA1-J-SC with anti-J chain clones followed by detection with anti-IgA. Representative plot of 2 experiments. E) SC binding. IgA1-J was captured with coated anti-J chain antibody, and then SC (site-specifically biotinylated via a C-terminal Bir A tag) was added. Representative plot of 2 experiments.

IgM is secreted as a pentamer consisting of five H2L2 subunits combined with one J chain, whereas incomplete polymers are ordinarily retained within the antibody secreting cell until properly assembled.^3^ In addition, IgM in circulation becomes complexed with a small scavenger molecule called CD5L in a covalent manner, making it an integral part of IgM’s composition.^4,6^

Although IgM is normally exclusively secreted as a J chain-containing pentamer, J chain-negative IgM can be found in some pathological conditions, which are assembled predominantly as hexamers.^4,5^ Evidence from *in vitro* culture systems, mainly using myeloma-derived IgM, and reports of aberrant IgM molecules in Waldenström Macroglobulinemia and cold agglutinin patients suggests that acquired mutations in B cells during lymphomagenesis may cause secretion of IgM without J chain. ^25,26^

IgA’s structure, on the other hand, is more varied. In addition to the existence of two different subclasses that mainly differ in hinge length (IgA1 > IgA2), IgA can be secreted as monomers and different types of polymers. Monomeric IgA is the result of IgA expression without J chain and makes up the bulk (about 80-90%) of all circulating IgA.^7,8^ Dimeric IgA is formed when J chain is co-expressed and is made up of two H2L2 subunits and one J chain. Low concentrations of higher order J chain-containing polymers have been reported as well, which can vary significantly on the clonal level.^8,9,57^ For IgA, studies indicate significant variability in the dynamics of monomeric and polymeric IgA responses in circulation to vaccines or infections.^27–31^ It appears that antigen-specific IgA responses may (temporarily) be dominated by dimeric IgA. Furthermore, we have recently observed that IgA autoantibody responses in several rheumatic diseases are comparatively skewed towards a high percentage of polymers.^49^ The role of dimeric IgA in acute and chronic (auto)immune responses is largely unknown; we lack insights into both the dynamics and functional consequences of polymeric antibodies.

Both IgM and IgA may be transported to mucosal sites via the polymeric Immunoglobulin receptor (pIgR), which binds in a J chain-dependent manner. Consequently, IgA at mucosal sites is (almost) exclusively polymeric.^10,11^ The pIgR shuttles bound polymeric antibodies directly to the extracellular, mucosal space where the receptor-antibody complex is subsequently cleaved of and released.^12,13^ The extracellular fraction of the pIgR remains attached to the polymeric antibody as secretory component (SC).^14^ In IgA, this interaction is covalent, whereas in IgM, binding is noncovalent. Secretory IgM is largely devoid of CD5L.^4^ In addition to pIgR, multiple other receptors show preferential binding to distinct polymeric forms of IgM or IgA. In particular, the Fcαμ receptor binds to both IgM and IgA and requires the J chain for its interaction with these ligands.^15,16^ Furthermore, FCRL3 and FCRL4 have been reported to selectively bind secretory IgA and circulatory polymeric IgA, respectively.^21,22^ While IgA does not activate complement, both IgM-J and IgM devoid of J chain is a much stronger activator of the classical pathway. ^23,24,58^ It is therefore relevant to be able to distinguish these structural IgM and IgA variants when monitoring immune responses.

Studies of polymeric composition of IgA and IgM in various immune settings are severely hampered by the lack of reliable, high-throughput assays. Typically, size-exclusion chromatography (SEC) or sucrose gradients have been used to separate polymers based on size,^27–31,50^ but these require a lot of hands-on work, limiting the number of study objects that can be included. Alternatively, IgA polymer contents may be assessed via the detection of J chain. Historic attempts relied on denaturation of IgA for J chain detection, since it was found that few epitopes on the J chain are available for binding in the intact IgA molecules.^51^ Antibodies that can reliably and quantitatively detect J chain as part of IgA (or IgM) do not appear to be commercially available. Several monoclonal antibodies that target integrated J chain in IgA have been reported, but without detailed evaluation it remains unclear how these perform with respect to detection of IgM vs IgA, IgA subclasses and allotypes, serum vs recombinant Ig, presence of CD5L or SC, etc.^32,52,53^ Detection of J chain with secretory component (or pIgR) appears to be the most feasible approach that has been attempted,^33,34^ but the relatively low affinity of the pIgR may limit assay development, for instance in the case of the detection of low-level, antigen-specific responses.

In this study, we report the generation of a novel panel of anti-J chain antibodies with proper binding characteristics for assessment of IgM- and IgA-integrated J chain. We have used well-defined IgM and IgA antibody preparations to extensively characterize these clones and have subsequently used them to setup assays that can reliably quantify integrated IgM-J and IgA-J, opening the door to extensive mapping of polymeric responses.

## Materials & methods

### Production of recombinant proteins

Recombinant IgM antibodies specific for citrullinated proteins (1G8 and 2D5) were produced using HEK293F and purified as described previously.^23^ Various IgA antibodies against SARS-CoV-2 spike protein (COVA1-18^35^) were produced similarly, either with or without co-transfection of a vector containing the coding regions for the J chain, in order to produce oligomeric IgA. Secretory IgA was made by additionally co-transfecting a vector coding for SC. We expressed two allotypic variants of IgA1 (IGHA1*01 and IGHA1*06, abbreviated as IgA1.1 and IgA1.6, resp.), and four allotypic variants of IgA2 (IGHA2*01, IGHA2*02, IGHA2*03, and IGHA2*05, abbr. similarly). IgA was purified using a peptide M Sepharose column (Invitrogen). In addition, human plasma-derived IgA was also purified using this column. In short, the filtered culture supernatant or plasma was loaded onto the column and the bound IgA was subsequently eluted with 0.1M Glycine combined with 0.5M Arginine (pH 3-3.5). The eluate was then neutralized with 2M Tris (pH 9.0) and a 10-kDa spin column (Amicon Ultra-4 Centrifugal Filter Unit; Merck) was used to concentrate and exchange the buffer to PBS. Antibodies were aliquoted and stored at -20°C.

### High-performance size-exclusion chromatography (HP-SEC) of antibodies

In order to check the quality of the produced antibodies and their polymerization state, antibody preparations were run with HP-SEC using a Superdex200 10/300 GL or Superose6 Increase 10/300 GL column (GE Healthcare) on an Agilent 1260 Infinity II HPLC system. 20 µg of each antibody was diluted in 120 µL PBS, loaded onto the column and eluted with PBS. The elution was monitored by measuring the absorption at 280 nm. To obtain separate fractions of plasma-derived IgA polymers and monomers, the purified plasma IgA was run similarly, but in larger volumes (100 µL serum + 200 µL PBS) and fractions of 250 µL were collected. The peaks corresponding to polymers and monomers were pooled and concentrated.

### Serum samples

Healthy donor sera were obtained from anonymous healthy volunteers with written informed consent, adhering to Dutch regulations and the Declaration of Helsinki. Peripheral blood serum samples of SARS-CoV-2 vaccinated healthy individuals were collected as part of the Target-2-B! cohort study upon written informed consent, approved by the medical ethical committee (NL74974.018.20 and EudraCT2021-001102-30).^54^

### Generation of monoclonal antibodies against J chain

Antibodies targeting human J chain were generated as described previously.^36^ Briefly, a rabbit was immunized and boosted every four weeks with recombinant human J chain produced in *E. coli* (LSBio). Blood was drawn after the 5^th^ booster, and PBMC’s were isolated. Through FACS sorting, cells were enriched for antigen-specific B cells based on staining for IgG and binding to labeled recombinant human J chain and recombinant IgM, seeded 1 cell/well, and cultured for 8-9 days. Culture supernatants were screened for antibodies specific for human J chain, IgA-J and/or IgM-J, and positive clones selected. From these clones RNA was isolated, cDNA was synthesized and IGHV and IGLV regions were amplified by PCR and then sequenced by Sanger sequencing. Antibodies were expressed using synthetic DNA vectors (GeneArt, Invitrogen) with mouse IgG1-kappa constant domains combined with the obtained variable regions and purified using a HiTrap protG column (GE Healthcare) as described previously ^23^. The purified IgG was stored in 5mM NaAc (pH 4.5) at -20°C.

### Determination of anti-J chain binding epitopes

The binding epitopes of the generated anti-J chain clones were tested in an enzyme-linked immunosorbent assay (ELISA). To map the binding of the different clones, Maxisorp plates (Thermo Fisher Scientific) were coated overnight with 1 µg/mL of anti-J chain antibody in PBS at 4°C. Plates were washed five times with PBS + 0.02% Tween-20 (PBS-T), and 0.1 µg/mL biotinylated human J chain was pre-incubated for 1h with different concentrations of the anti-J chain antibodies in PBS supplemented with 0.2% w/v gelatin (Merck) and 0.1% v/v Tween-20 (PTG), with the highest concentration of 2 µg/mL. Then 100 μL of this mixture was added to coated plates and incubated for 1h at RT while shaking. Plates were washed five times with PBS-T, and bound J chain was detected with HRP-labeled streptavidin (strep-HRP, 1:1000) in PTG for 30 min. The plates were washed again and binding was visualized with 100 μl 3,3′,5,5′-tetramethylbenzidine (100 μg/mL) in 0.11 M acetate buffer (pH 5.5), supplemented with 0.003% H_2_O_2_ (Merck). The reaction was stopped with 100 μl of 0.2 M H_2_SO_4_ and the plates were read using a Biotek plate reader. The level of inhibition was determined for each anti-J chain coat as OD (inhibition condition) / OD (no inhibition) * 100%. Reactivity towards integrated J chain was tested by incubating varying concentrations of recombinant IgA1.1 (anti-spike) and IgM (ACPA 2D5) produced with and without J chain on plates coated with either mouse-anti-human IgA (MH14-1, Sanquin) or mouse-anti-human IgM (MH15-1, Sanquin). Bound IgA and IgM were detected using biotinylated anti-J chain clones, followed by incubation with Strep-HRP.

### Development of an IgM-J assay

We continued with the development of an IgM-J ELISA that would work in full serum and therefore focused on clones 1G5 and 8H4, which showed the most favorable binding profile towards IgM and limited binding to IgA. Plates were coated overnight with mouse-anti-human IgM (1 µg/mL in PBS, MH15-1, Sanquin) at 4°C. Recombinant IgM (ACPA 2D5) with and without J chain, and a reference serum pool with a known concentration of IgM were tested in various concentrations in PTG. Alternatively, the recombinant IgM-CD5L complexes were tested on an antigen coat, for which a biotinylated citrullinated cyclic peptide (CCP4-bt; 2 μg/mL in PBS-T) was allowed to bind to streptavidin coated plates (Thermo Fischer Scientific). Integrated J chain was detected with biotinylated anti-J chain clones (1 µg/mL) and subsequently with streptavidin-poly-HRP (1:5000; Sanquin). As binding of the anti-J chain clones was impaired to serum-derived IgM, likely due to the presence of CD5L on IgM from serum, we tried selective removal of CD5L using mildly reducing conditions as we have done previously^4^. We tested several conditions, i.e. different concentrations (1 to 3mM) of glutathione (GSH, Sigma), and incubation times (2h to 22h), and the ability of both a-JC clones (1G5 and 8H4) to detect serum-derived material. For the final assay, serum samples were prediluted in tris-buffered saline (TBS) supplemented with 0.1% Tween-20 (TBS-T), 1 mM GSH and 10 mM Ethylenediaminetetraacetic acid (EDTA), and incubated overnight at 37°C while shaking at 300 rpm. After this incubation time, the samples were further diluted in TBS-T and added to plates coated with anti-IgM (MH-15-1). Recombinant IgM-J was included as a calibrator. Integrated J chain was detected with HRP-labeled anti-J chain 1G5 in TBS-T (0.25 μg/mL) and visualized as described earlier.

### Detection of IgA-J

To detect IgA-J we decided to continue with clone 9G10, as it appeared to be the only clone that could properly bind to serum-derived IgA (**Figure S4**). Plates were coated overnight with anti-IgA (1 μg/mL, MH-14-1) in PBS at 4°C. Recombinant IgA1.6 (anti-spike), purified serum-derived monomeric IgA (mIgA) and polymeric IgA (dIgA) and serum samples were diluted in PTG and 100 μl was added to the plates after washing. Integrated J chain was detected with biotinylated anti-J chain 9G10 and subsequently with streptavidin-poly-HRP (1:10.000). A serum pool with a known concentration of IgA and recombinant IgA-J were taken along as calibrators. Isotype-specific J chain-detection followed a similar protocol, but instead anti-J chain 9G10 was coated (1 μg/mL) and bound IgA was detected with HRP-labeled mouse-anti-human IgA1 (1 μg/mL, MH141-1-HRP, Sanquin) for IgA1 and mouse-anti-human IgA2 (1:2000, A9604D2-HRP, Southern Biotech) for IgA2 in PTG. IgA1.6-J (anti-spike) and IgA2.1-J (anti-spike) were taken along as calibrators. Total IgA1 and IgA2 levels were determined analogously, in assays using anti-IgA coat as capture step.

### Detection of antigen-specific IgA-J

Maxisorp plates were coated with recombinant SARS-CoV-2 spike protein (1 μg/mL) at 4°C overnight. Samples were diluted in PTG. Additionally, recombinant IgA1.6-J (anti-spike) was taken along as a calibrator. Coated plates were washed, 100 μl of sample dilution was transferred and incubated for 1 h at room temperature. IgA-J was then detected with biotinylated anti-J chain 9G10 (0.25 μg/mL, 1 h incubation) and subsequently with streptavidin-poly-HRP (1:10.000, 30 min incubation). The reaction was visualized and plates were read.

### IgM and IgA depletion

For some experiments, samples were depleted of IgM or IgA. Serum samples were diluted 30-100x in PBS-T containing 10-20 mg/mL CaptureSelect IgM-XL or IgA-XL Sepharose beads (ThermoFisher) and were transferred to a 96-well polypropylene filter microplate (0.45 µm polyvinylidene fluoride membrane, Agilent). These plates were incubated overnight at room temperature while shaking at 300 rpm. The next day, the filter plates were centrifuged for 1 min at 500g, filtrates were collected in a 96-well microtiter plate. Depletion efficiency was examined by ELISA and found to be >95%.

### SPR

Surface plasmon resonance (SPR) measurements were carried out on an IBIS MX96 (IBIS technologies) as described before.^55^ Briefly, 9G10 was expressed as recombinant Fab with C-terninal Bir-A tag, biotinylated, and captured onto a SensEye G-streptavidin sensor (Ssens, Enschede, Netherlands) at 1 – 30 nM. IgA (anti-spike IgA1.6-J was injected at two-fold dilutions ranging from 5 down to 0.15 μg/mL. Sensorgrams were analyzed with Scrubber software version 2 (Biologic Software, Campbell, ACT, Australia).

### Data analysis and visualization

Graphs and statistical analyses were made using Graphpad Prism v10.

### Data availability statement

The data that support the findings of this study are available from the corresponding author upon reasonable request.

## Results

### Generation of monoclonal antibodies against J chain

In order to generate novel monoclonal antibodies against J chain, we immunized a rabbit with recombinant human J chain, and screened for and isolated several clones with promising anti-J chain activity (**Figure S1A-B**). After screening, we continued with eight clones that were able to bind to J chain in IgM-J, IgA-J, or both, as well as one clone that essentially only bound free J chain (**Figure S1C**). These clones were then subjected to further characterization.

### Anti-J chain mAbs show differential binding to IgA-J and IgM-J

To gain insight into the binding characteristics of the generated anti-J chain mAbs, we first tested if they compete with each other (**Figure 1B**). This revealed four clusters of anti-J chain mAbs that bind to distinct areas on the J chain. Cluster 1 consisted of 6F10, 9G10, 18D2 and 19E3, cluster 2 of 2D3 and 2G7, cluster 3 of 8H4 and 1G5 and then lastly, cluster 4 of 17H5. Next, we compared relative binding of the clones to recombinant IgA and IgM antibodies, with and without J chain (**Figure 1C, Figure S2A-B**). In all cases, binding to IgM and/or IgA with J chain was observed, whereas the control antibodies without J chain did not show appreciable binding. Between the abovementioned clusters, notable differences in preference for IgA-J vs IgM-J were observed. Clones within cluster 1 showed up to hundred-fold preferential binding to IgA-J over IgM-J. Furthermore, clone 9G10 bound much more avidly in comparison to the other clones. Cluster 2 showed approximately ten-fold preferential binding to IgA over IgM. Binding to IgA and IgM was broadly similar in cluster 3, whereas the sole clone in cluster 4 bound very poorly to integrated J chain, as expected based on the initial screenings. These binding patterns were similar for all IgA subclasses and allotypes (**Figure S2C)**. We also tested whether association of IgA with secretory component (SC) interferes with binding of the anti-J chain mAbs. In a direct capture assay, only clone 9G10 was observed to bind IgA-SC with similar efficiency as IgA-J without SC (i.e., IgA-J; **Figure 1D**). Furthermore, free SC was able to bind to IgA-J captured by clone 9G10 (i.e., SC and 9G10 can simultaneously bind IgA-J), whereas this was not the case for any of the other ant-J chain clones (**Figure 1E**).

### Successful quantification of IgM-J by immunoassay upon selective CD5L dissociation

For detection of J chain in IgM, we continued with the mAbs from cluster 3 (8H4 and 1G5) to develop an IgM-J chain assay, as they showed the most favorable binding profile to this end. We first compared our results with the recombinantly produced IgMs (with and without J chain) to serum IgM and found that the signal for the serum-derived IgM was lower than what we found for the recombinantly produced IgM-J (**Figure 2A**). This is likely due to the presence of CD5L in serum IgM, which is saturated with CD5L.^4^ Indeed, recombinant IgM-J-CD5L complexes showed a similarly reduced signal compared to IgM-J as serum IgM (**Figure 2B**). We remedied this by testing several conditions for selective CD5L removal from serum IgM to allow for J chain detection. A very mild reduction step was found to suffice to allow quantitative recovery of IgM-J from serum IgM (M&M, **Figure S3, Figure 2C**). We applied this assay setup to 20 healthy donor sera (**Figure 2D**) and found as expected that the amount of IgM-J chain is the same as the amount of total IgM (*r* = 0,99, p <0.0001), in line with our earlier finding that all circulating, normal IgM contains J chain _4_.

**Figure 2.**
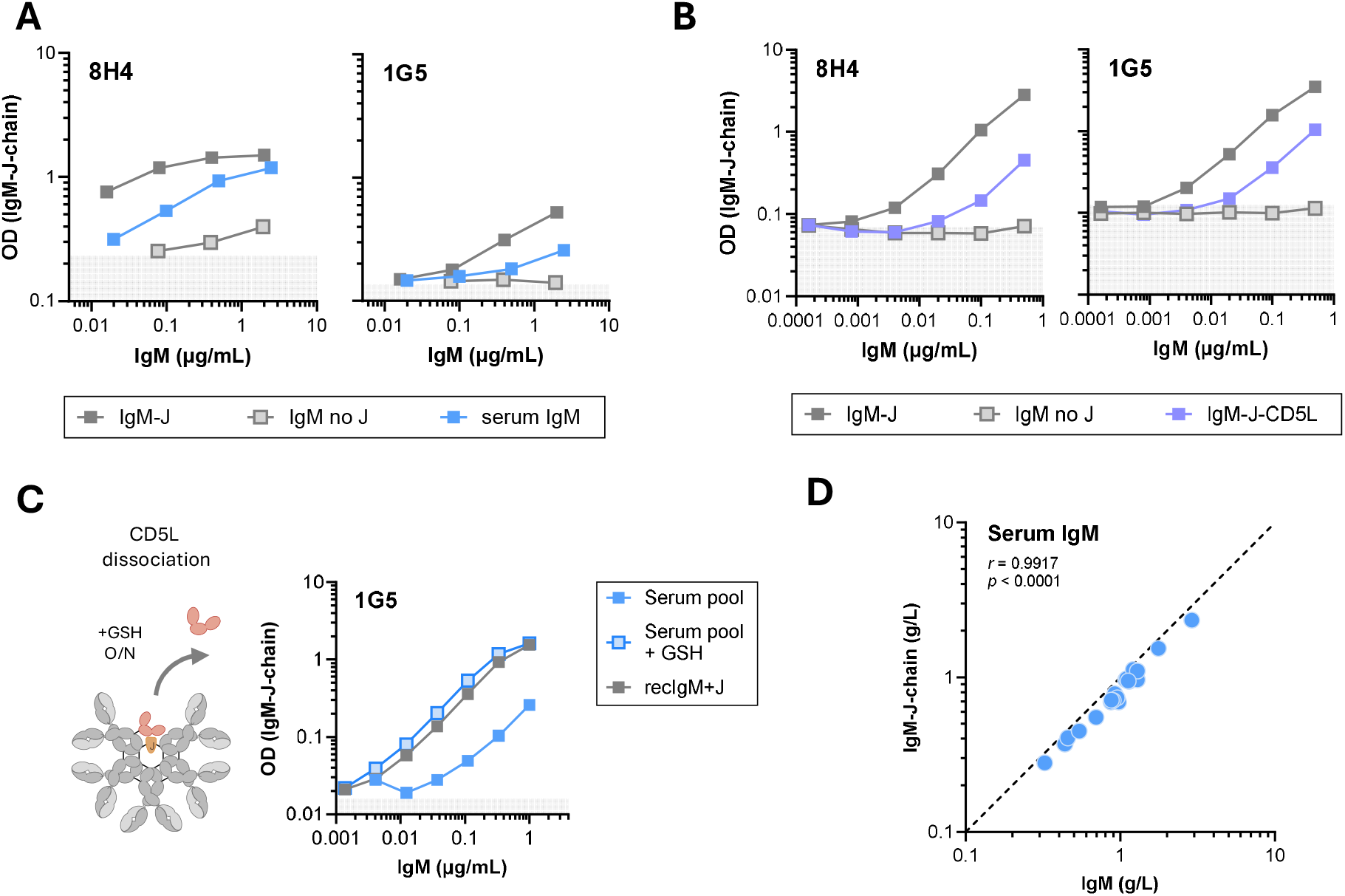
Development of IgM-J-chain assay. A) Binding of the anti-J chain clones 8H4 and 1G5 to recombinant IgM with and without J chain and serum-derived IgM, and B) to recombinant IgM-CD5L complexes. C) CD5L was selectively dissociated from serum-derived IgM via mild reduction with glutathione (GSH). Conditions were chosen for optimal CD5L removal, while leaving the IgM molecule intact (Figure S3). Anti-J chain clone 1G5 was the most promising in our tests and was then used with to detect IgM in a normal serum pool and a GSH-treated serum pool. A-C representative plots of 2-3 experiments. D) Detection of total IgM and IgM-J levels in n=20 healthy donor sera. Each dot represents 1 donor

### Development of an IgA-J chain ELISA

For the detection of IgA-J we continued with the clones from clusters 1 and 2, that all showed preferential binding to integrated J chain in IgA over IgM. All clones were able to bind to recombinant IgA-J as well as IgA polymers purified from plasma (**Figure 3A; Figure S4, S5A**). However, upon assessment of IgA directly from plasma, binding of most clones was observed to be substantially less than expected based on the ~10% IgA polymers found in circulating IgA on average.^37,50^ Only clone 9G10 appeared to be able to bind to both purified and non-purified IgA-J from plasma in a consistent manner and therefore was used for the continuation of assay development. We compared the performance of 9G10 in detecting IgA-J for plasma IgA and the purified polymeric and monomeric IgA pools from plasma of six donors (**Figure 3B**). Detected IgA-J and total IgA concentration of the polymeric fraction correlated well (*r* = 0.84; p < 0.05), showing that we can properly quantify IgA-J when it is purified. Similarly, the amount of IgA-J detected in plasma is around the expected value of ca. ~10%.^37,50^ Residual binding to the monomeric pool is <1%. We also fractionated the plasma of four donors using HP-SEC and assessed individual fractions for both total IgA and IgA-J content. With IgA-J detection, a single peak of polymeric IgA was observed, without any monomeric IgA detected (**Figure 3C; Figure S5B**). Combined, this shows that 9G10 binds to only IgA-J in this setup and our assay can quantify both recombinant and plasma-derived IgA-J reliably. To gain insight in overall J chain distribution among IgA subclasses, we measured total IgA-J and subclass-specific IgA-J in 64 healthy donors (**Figure 3D**). We found that IgA-J makes up just under 10% of circulating IgA. Interestingly, we found similar results for both IgA1-J and IgA2-J of 11% and 10%, respectively. We also tested the ability to reliably measure secretory IgA. IgA and IgA-J was measured in saliva of healthy individuals, which are expected to only contain secretory IgA, which contains the J chain. IgA-J measurements corresponded closely to total IgA (**Figure 3E**).

**Figure 3.**
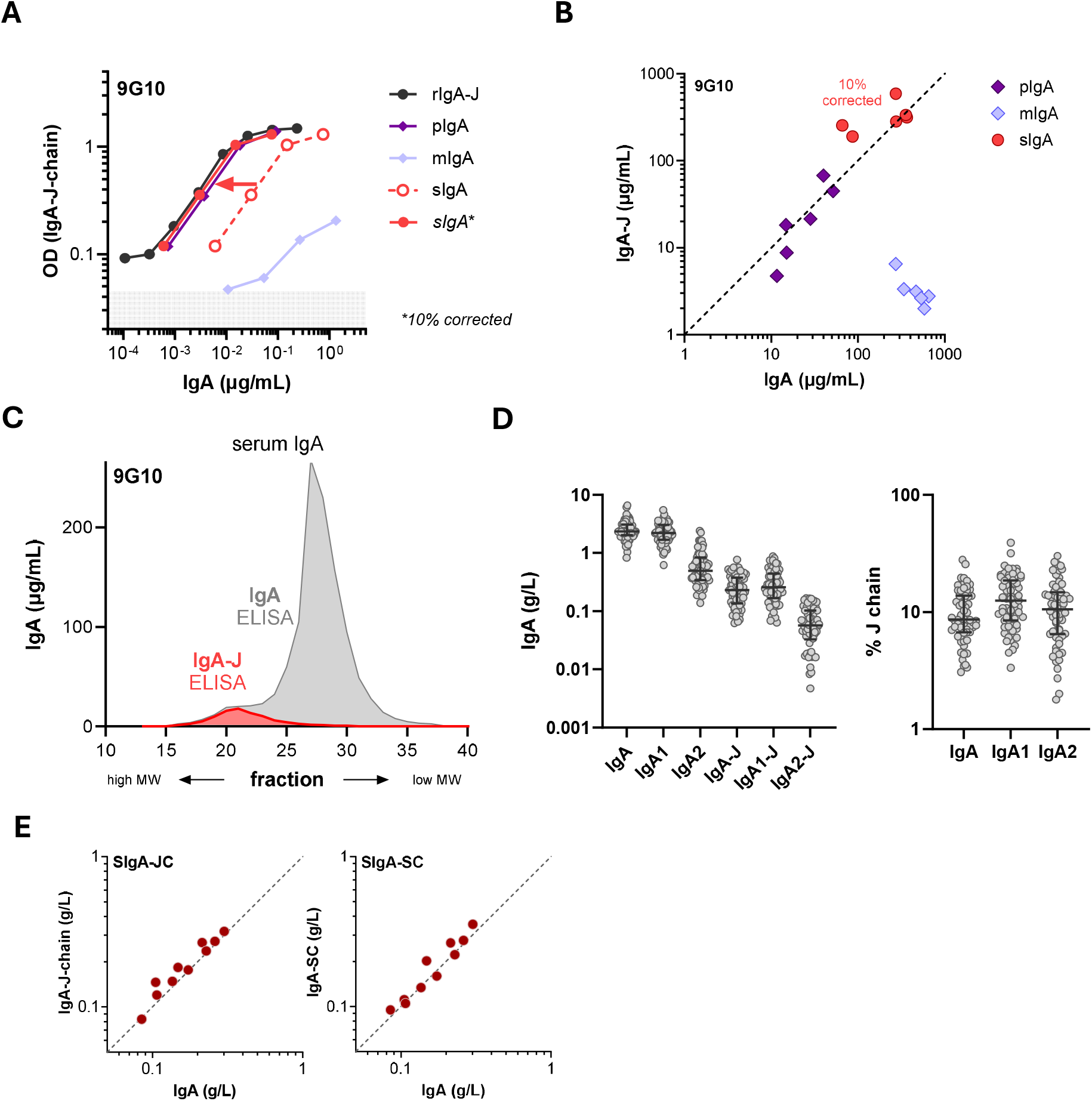
Development of IgA-J-chain assay. A) Detection of IgA-J chain with clone 9G10 of recombinant IgA1.6-J anti-spike 1-18 (rIgA-J), serum IgA (sIgA), and IgA, affinity-purified from serum and separated by HP-SEC into monomeric and polymeric IgA fractions (resp. mIgA and pIgA) (Suppl. Fig. 4). *sIgA also plotted vs estimated pIgA concentration of ca. 10% of sIgA, see main text. B) IgA-J vs IgA ELISA results for n=6 donors (each dot is one donor), for mIgA, pIgA, and sIgA (sIgA plotted vs estimated pIgA concentration of ca. 10%). IgA1.6-J was used as calibrator. C) Detection of IgA and IgA-J in fractionated serum from a healthy donor (representative of n=4; also see Figure S5B). D) Detection of IgA, IgA1, and IgA2 as well as IgA-J, IgA1-J, and IgA2-J in a panel of healthy donors (n=64). Each dot represents one donor. Right panel shows respective calculated percentages for J chain-containing IgA, IgA1, and IgA2. E) Detection of IgA-JC and IgA-SC in saliva of 10 healthy individuals. Each dot represents a single individual.

### Quantification of antigen-specific IgA-J

Next, we explored the possibilities for detecting IgA-J in an antigen-specific manner, which will be useful for studying the variable dynamics of polymeric IgA during various immune responses. To this end, we focused on the antibody response against SARS-CoV-2, for which we previously demonstrated variable and substantial contributions of polymeric IgA.^49^ Clone 9G10 was found to have a fairly high affinity of ca. 0.5 nM (**Figure S6**), and is therefore expected to be suitable for generating sensitive immunoassays. In addition, this antibody has a strong, ca. 100-fold preference for binding IgA-J over IgM-J, both for IgA1 and IgA2 (**Figure 4A**), suggesting the possibility for selective IgA-J detection in immunoassays. IgM-J binding was not further impacted by the presence of CD5L (**Figure S7**). We analyzed a panel of serum samples from healthy individuals vaccinated against SARS-CoV-2 by capturing antibodies to immobilized spike protein followed by detection of J chain with 9G10. Whereas a clear correlation with IgA anti-spike was apparent, no correlation was observed with IgM anti-spike present in these serum samples (**Figure 4B**). Samples were also tested after depletion of total IgM or IgA. Whereas depletion of IgM does not have an appreciable impact on either IgA anti-spike or anti-spike antibodies detected via J chain, IgA depletion results in a large reduction in the J chain signal (**Figure 4C**). In a previous study^49^ we fractionated post-vaccination sera using HP-SEC to analyze anti-spike polymeric IgA. Here we analyzed elution profiles for anti-spike antibodies containing J chain in three of these samples. This resulted in elution profiles coinciding with the high-molecular weight IgA part of the anti-spike IgA elution profile (**Figure 4C**). We also analyzed the serum samples from this study directly to estimate the fraction of IgA-J anti-spike. Anti-spike IgA-J and IgA levels were measured, and we compared the percentage of IgA-J obtained via SEC analysis (determination of the area under curve of both peaks) with the values obtained via ELISA (**Figure 4D**). These compare well (*r* = 0.78; p < 0.0001), showing that the new assay is able to properly quantify the antigen-specific polymeric IgA responses to SARS-CoV-2. Taken together we show that clone 9G10 specifically detects IgA-J, enabling us to reliably quantify antigen-specific polymeric IgA in a high-throughput manner.

**Figure 4.**
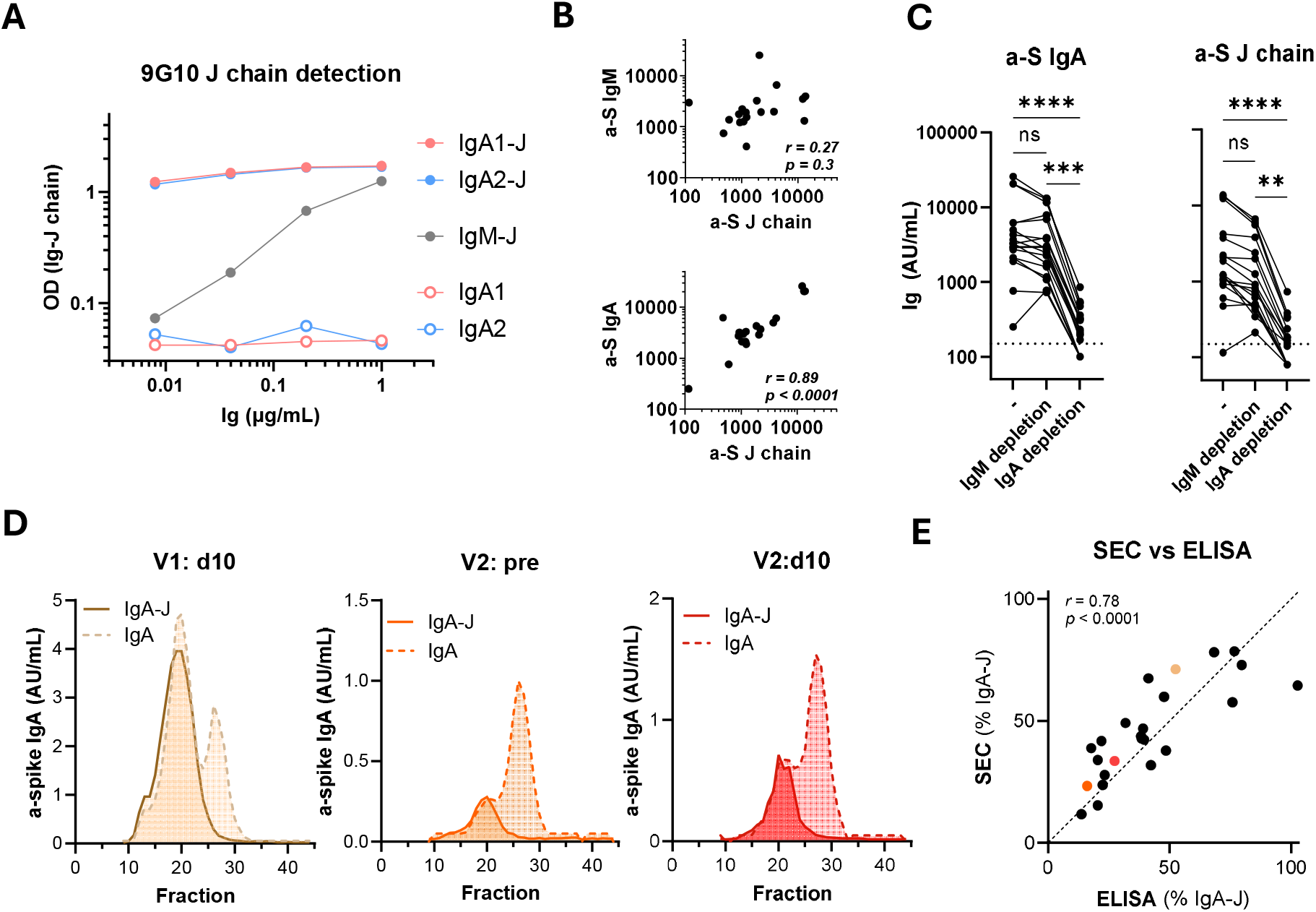
Development of antigen specific IgA-J chain assay. A) Comparison of J chain detection using clone 9G10 for recombinant IgA1, IgA2, and IgM (anti-spike clone 1-18). B,C) Detection of anti-spike (a-S) IgA, IgM, or J chain in serum samples of healthy individuals (n=19) vaccinated against Sars-CoV-2. Each dot represents one individual. B) Comparisons between J chain detection and IgM or IgA detection are shown on top and bottom respectively. Pearson correlation coeficient is indicated as *r*. C) J chain detection upon depletion for either IgM or IgA, in comparison to non-depleted (-). Friedman test with Dunn’s multiple comparisons; ns not significant, ** *p*<0.01, *** *p*<0.001, **** *p*<0.0001. D) Sera of a healthy donor after SARS-CoV-2 vaccination at three time points (day 10 after first injection ‘V1 d10’, and just prior to and 10 days after second injection, ‘V2: pre’ and ‘V2: d10’, resp.), were fractionated using HP-SEC, fractions were analyzed with IgA anti-spike ELISA and IgA-J chain anti-spike ELISA (9G10 detection). E) Comparison of % IgA-J determined as AUC of the polymer and monomer peaks of HP-SEC elution profiles, vs % IgA-J obtained with ELISA in 24 sera from 8 healthy donors. Pearson correlation coefficient is indicated as *r*.

## Discussion

Here we describe the generation and characterization of novel monoclonal antibodies that are able to detect J chain in IgA and IgM. We show that we can use these clones to reliably detect and quantify IgM-J and IgA-J, and that we can monitor antigen-specific polymeric IgA responses over time. These assays allow for high-throughput mapping and more accurate quantification of these responses than for example the use of SDS page or HP-SEC. Using these methods, we were able to readily demonstrate the polymer content of circulatory IgA1 and IgA2, which are remarkably similar (both around 10%), and demonstrated the feasibility of monitoring the dynamics of circulatory IgA-J during a vaccination response.

We have generated several different anti-J chain clones binding to what appears to be four different areas on the J chain. After thorough testing we found that they bound IgM-J and IgA-J with variable preference, a feature we used to further develop assays to detect both separately and specifically. Most clones appear to preferentially bind to IgA-J over IgM-J, which may be explained by slight structural differences described in J chain folding in both polymers.^38,39^ Alternatively, the additional tailpieces in an IgM pentamer might cause some steric hinderance.^39^ Lastly, we found discrepancies in binding to recombinant and serum-derived IgA-J and IgM-J. For IgM-J this could be pinpointed to the presence or absence of CD5L, which could be remedied by selective removal of CD5L for serum IgM using a simple mild reduction step. For IgA-J, we have not fully detailed where these discrepancies originate from. Possibly, another interfering factor exists, that is seemingly lost after purification. Regardless, these discrepancies were not observed with clone 9G10, which is able to bind similarly to both recombinant and serum-derived preparations in a consistent manner. All in all, for IgM, we have established a robust protocol for J chain detection using selective CD5L dissociation and detection with a clone that shows good binding to IgM-J (1D5). For IgA, the 9G10 clone in particular has good overall properties for J chain detection, i.e., high affinity, no interference by SC, no apparent preference for IgA1 vs IgA2, nor serum-derived IgA vs recombinant IgA, and a strong (~100-fold) preference for IgA-J over IgM-J.

For serum IgM we find, as we also showed previously ^4^, that all IgM in circulation exists as a J chain-containing polymer. This again suggests that the notion that part of normal human IgM is hexameric, is incorrect. However, multiple patients with B cell-driven disorders have been described that show reduced or even absent integration of J chain. We and others have shown that IgM can be produced in absence of J chain in certain conditions, such as Waldenström’s macroglobulinemia and cold agglutinin disease.^25,26,56^ In particular, we have applied the method described in this study to a pilot cohort of patients with Waldenström’s Macroglobulinemia and found that approximately one third of patients have IgM that is at least partly J chain-deficient.^56^ It is currently unclear what the implications of J chain-negative IgM are for the course of these diseases or if there even are any. The issue of most concern is the potential for a more pathogenic IgM (auto)antibody that activates complement way more efficiently than its “normal”, pentameric counterpart.^23,40^Larger patient cohorts need to be studied to truly answer this question. Furthermore, it is likely that there are other B cell-driven diseases that may also present with similarly defective IgMs, which is a matter to be explored further.

Because of the co-occurrence of monomeric and polymeric IgA in the human body, the study of IgA polymerization is more difficult than that of IgM. Reports on the relative contribution of IgA1 and IgA2 fraction, but also the dimeric and monomeric fraction, to total circulating IgA vary, especially for the latter. We find that polymeric IgA makes up approximately 10% of total serum IgA as reported by some earlier,^37,50^ but it ranges from 5 to 30% between donors, showing large variation between individuals. Interestingly, we observed similar percentages of J chain-containing IgA for both circulating IgA1 and IgA2, contrary to the implicit notion that a larger fraction of IgA2 is polymeric. It seems that IgA2 is not overrepresented in the polymeric fraction, despite its connection to mucosal immunity, and that some other factor drives J chain integration.

It is currently not completely understood how B cells decide to develop into cells that secrete IgA without J chain. Either J chain is downregulated again during differentiation/proliferation or the protein is degraded *via* another pathway in these cells.^42^ It is similarly unclear what events lead to the formation of J chain-negative IgM in some patients with B cell malignancies. The regulation of J chain expression occurs at the level of B cell maturation by master regulator PAX5,^43^ which is downregulated during differentiation of B cells into antibody-secreting cells.^44^ As PAX5 inhibits expression of J chain, this means that it is universally upregulated in antibody-secreting cells regardless of its isotype. Indeed, IgG B cells in human tonsils also stain positive for J chain.^45^ Studying the polymeric dynamics of different responses may provide indirect clues as to the conditions that help decide the fate of these J chain-positive B cells.

In conclusion, we have generated novel monoclonal antibodies that can detect integrated J chain in both IgA and IgM and used these to set up assays that can detect polymeric antibody responses with ease and accuracy. This opens the door to further investigation of aberrant IgM molecules in B cell driven diseases and of normal and pathogenic IgA responses and dynamics.

## Acknowledgment

We would like to thank Arthur Bentlage for the affinity measurements and Cristina Rada Santacruz for help with the experiments. This work was supported by a ZonMw grant (10430072010007).

