## Supplemental Information for "Assessment of polymeric immunoglobulin A and M via detection of the joining chain"

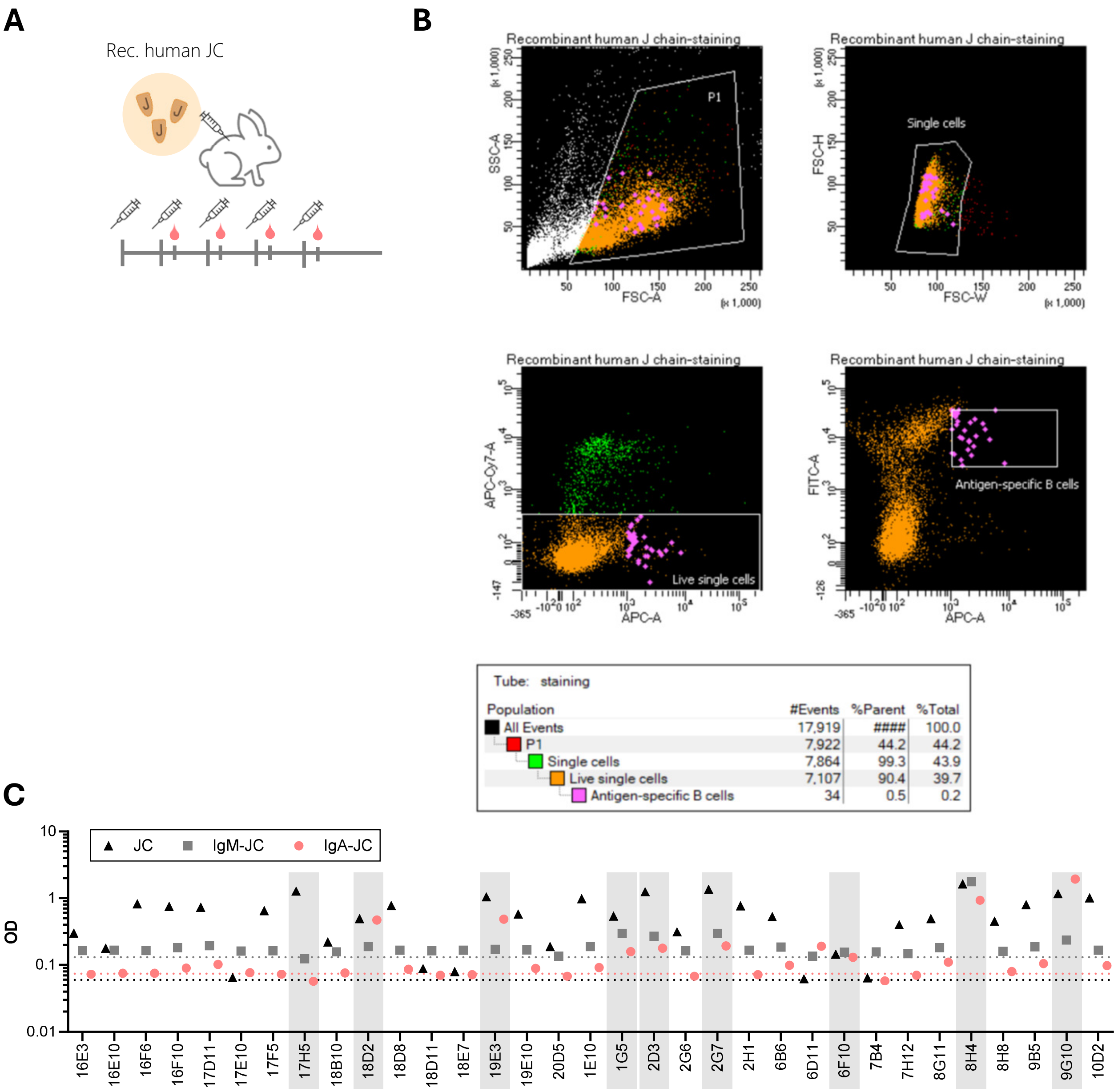


**Figure S1 | Screening of B cell supernatants for anti-J chain antibodies**

A) Immunization of rabbit with recombinant human J chain. Monthly injections were followed by blood draws 9 days later.

B) Sorting strategy for enriching for J chain-specific single B cells

C) Initial screening of culture supernatants of sorted single B cells, after 8-9 days of culture (see Materials and Methods for details). Shown are results of clones positive for anti-J chain, results in IgM-J and IgA-J screening assays, and selection of clones (highlighted in gray) for recombinant production and further analysis.

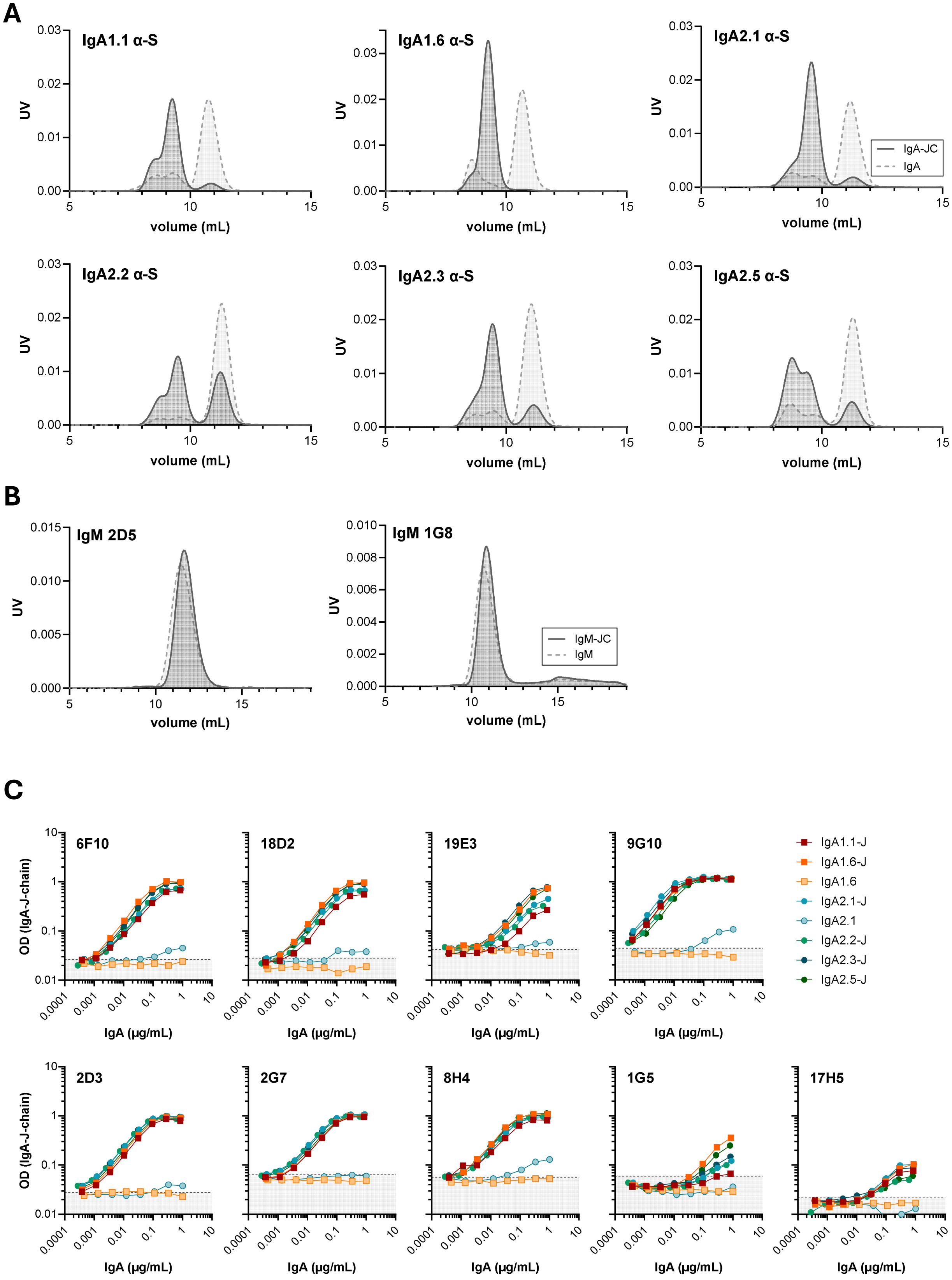


**Figure S2 | Characterization of recombinant IgA and IgM with and without J chain**

A) HP-SEC analysis of the anti-spike IgA antibodies used. Anti-spike clone 1-18 was expressed as different IgA subclasses and allotypes with or without J chain. The numbers refer to the IMGT numbering: e.g. IgA1.1 corresponds to IGHA1*01, etc.

B) HP-SEC analysis of IgM antibodies used (ACPA clone 2D5 and 1G8).

C) Binding of all anti-J chain clones against IgA-J subclasses and allotypes and IgM

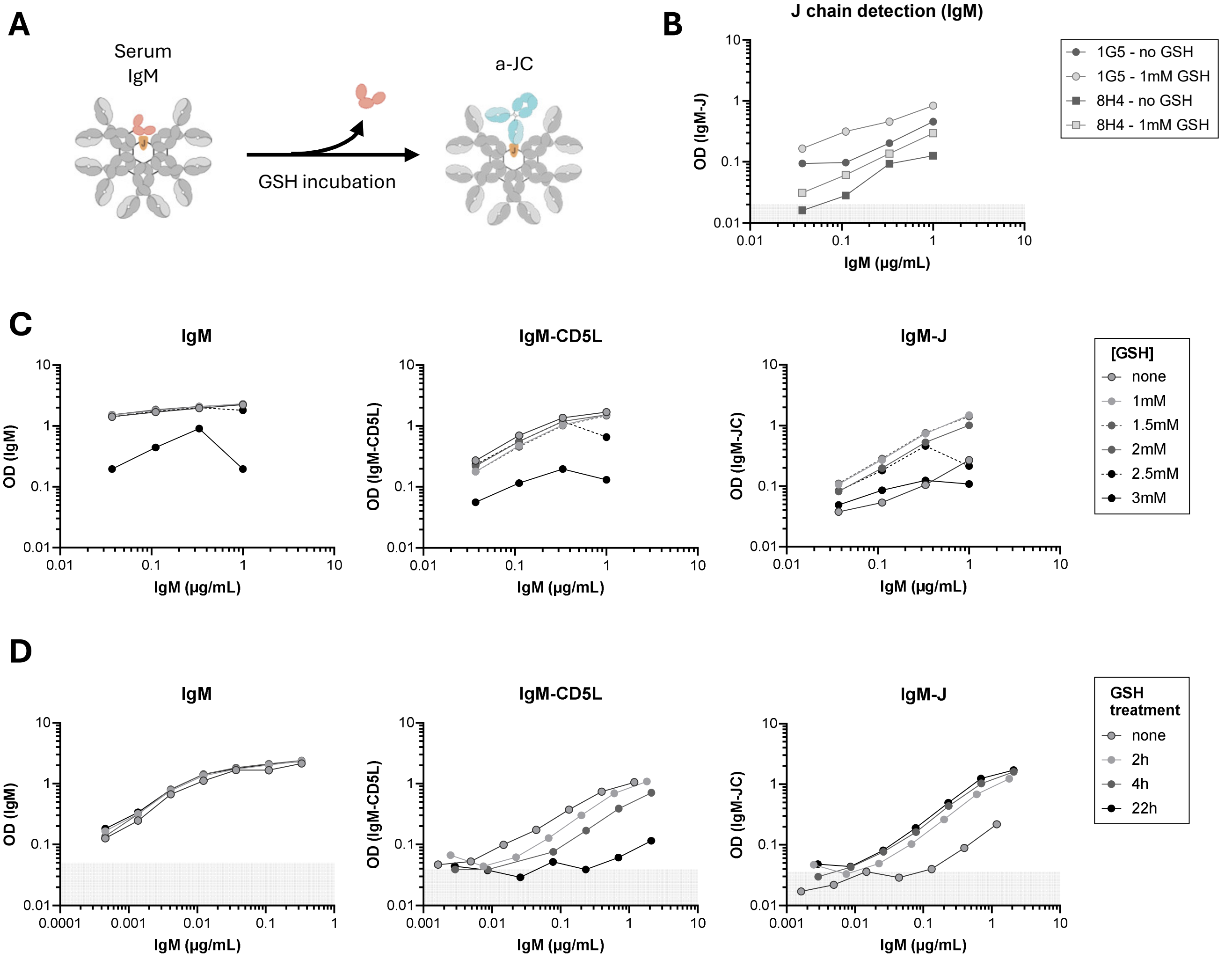


**Figure S3 | Optimization of IgM-J assay**

A) Schematic of GSH incubation of serum IgM

B) Comparison of 1G5 and 8H4 anti-J chain antibodies on GSH-treated serum pool.

C) A serum pool with a known concentration of IgM was incubated with varying concentrations of GSH for 2.5h at 37 °C. 1 mM of GSH results in effective J chain detection.

D) The same serum pool was incubated with 1mM GSH for 2h, 4h and 22h. After 22h most of the CD5L has dissociated, resulting in the best IgM-JC detection.

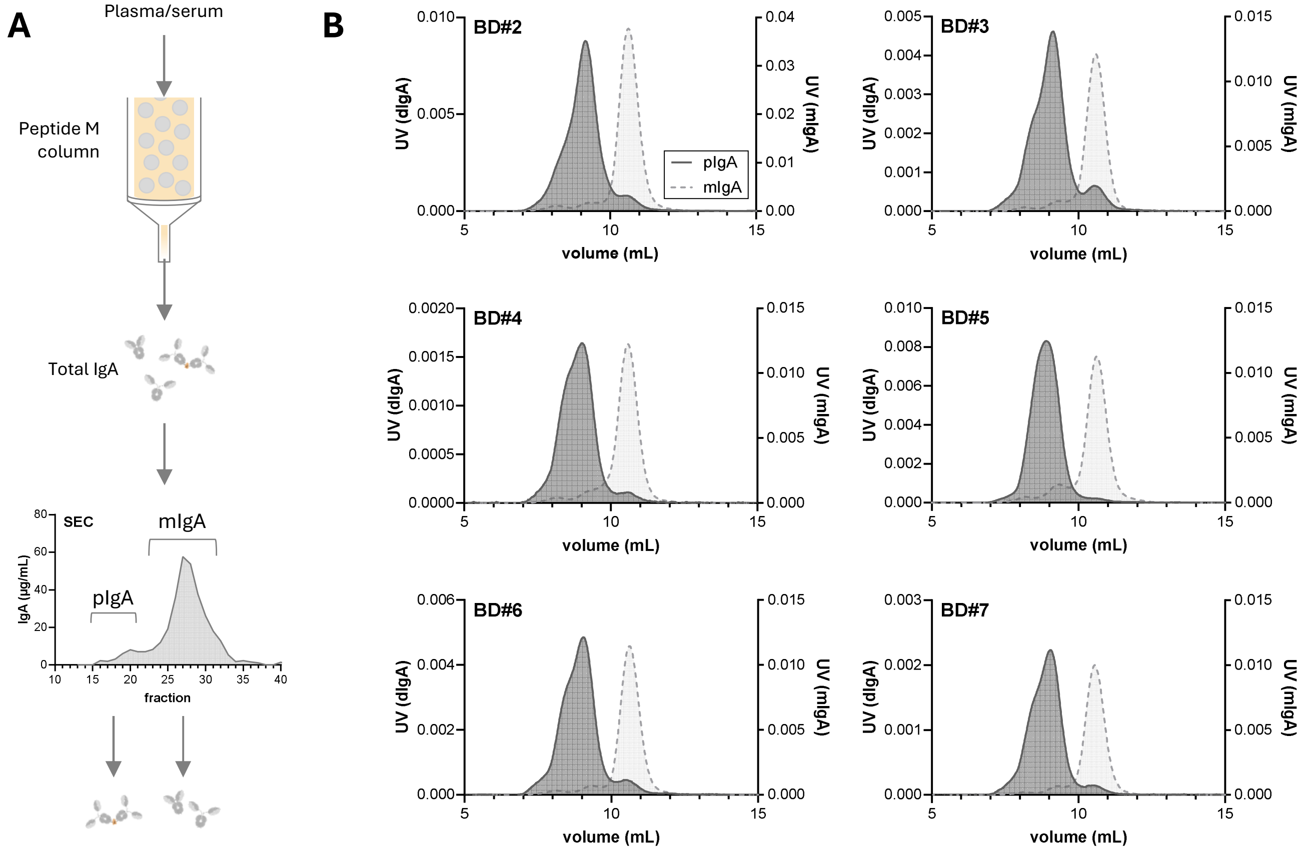


**Figure S4 | Purification of pIgA and mIgA from plasma or serum**

A) Schematic overview of method of purification. First step is affinity capture using peptide M resin. Second step is HP-SEC separation by size.

B) SEC of purified, pooled pIgA and mIgA preparations from plasma or serum

**A**

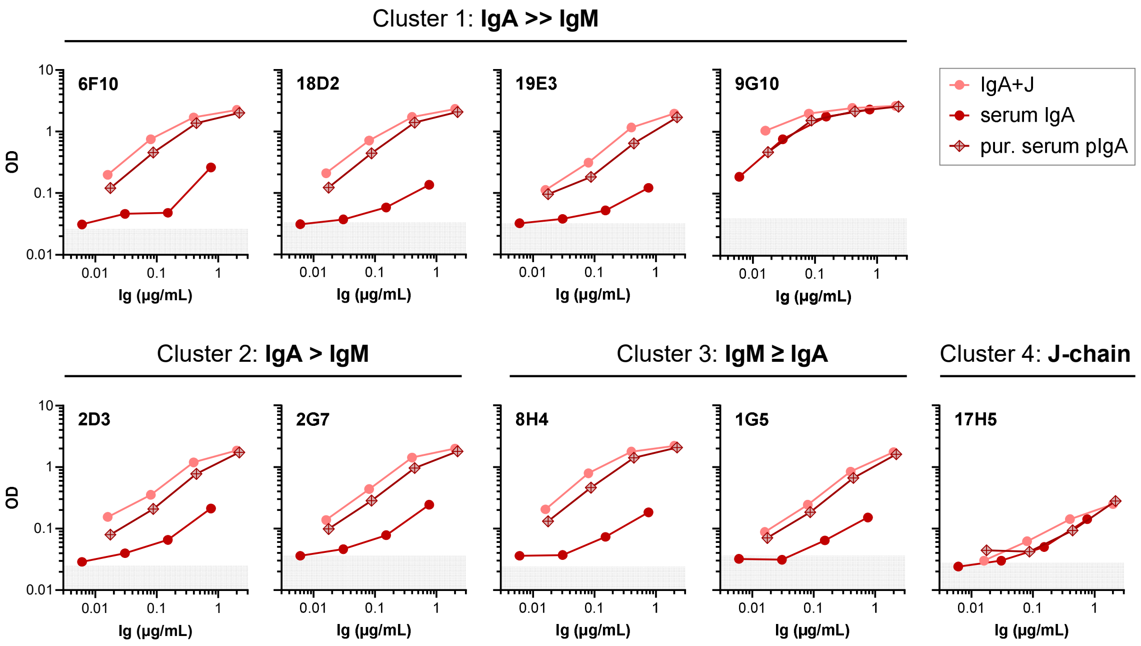


**B**

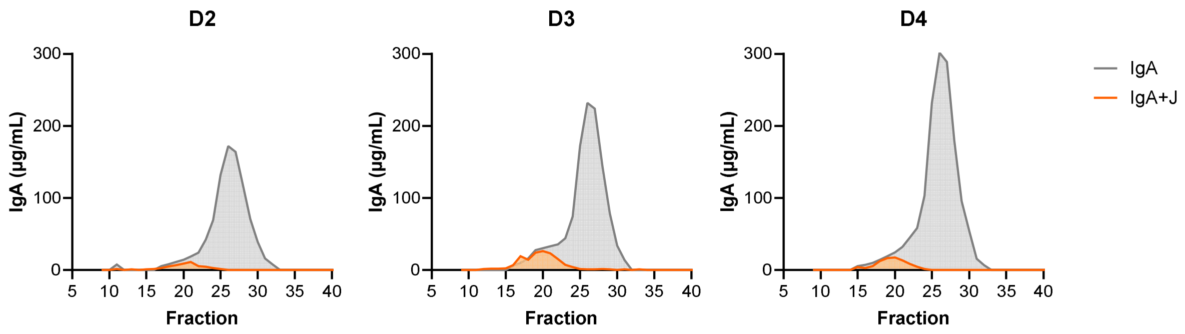


**Figure S5 | Detection of J chain IgA in serum or IgA purified from serum using different anti-J chain antibodies**

A) Capture of IgA using coated anti-IgA antibody, followed by detection using one of the anti-J chain clones. Tested are both serum and purified serum polymeric IgA, in comparison to recombinant IgA-J (anti-spike clone 1-18 IgA1.6-J). For serum, data is plotted against an estimated IgA polymer content of 10%. For most clones, but not 9G10, detection of polymeric IgA directly from serum apears less efficient then for purified serum polymeric IgA.

B) Detection of IgA and IgA-J in fractionated healthy donor sera.

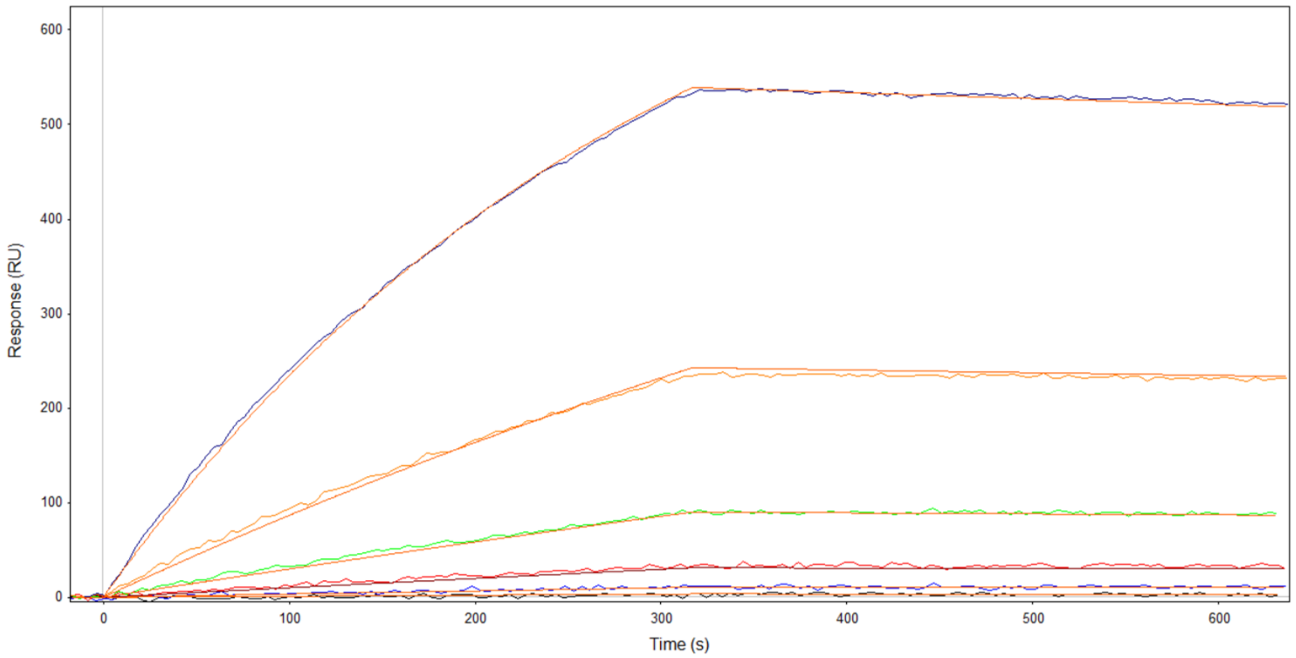


**Figure S6 | Affinity of 9G10 determined by SPR.**
A Fab fragment of 9G10 with an N-terminal BirA tag was site-specifically biotinylated and captured onto a streptavidin sensor. IgA1.6-J was injected at concentrations between 0.15 – 5 µg/mL. $k_a$ = 1.9×10$^5$M$^{-1}$s$^{-1}$, $k_d$ = 1.2×10$^{-4}$ s$^{-1}$, $K_d$ = 5×10$^{-10}$M.

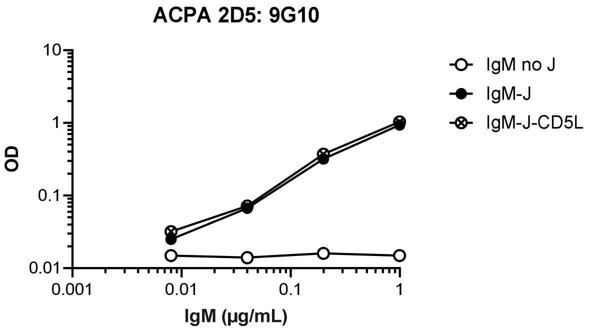


**Figure S7 | impact CD5L on 9G10 binding.**
Binding of the anti-J chain clone 9G10 to recombinant IgM with and without J chain and with CD5L.

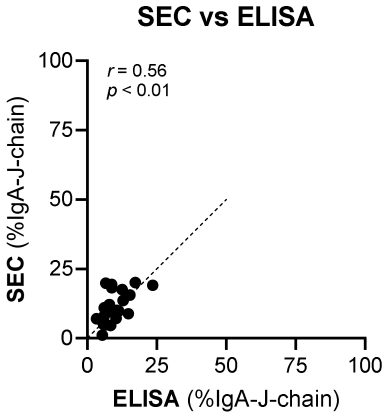


**Figure S8 | HP-SEC vs elisa for total serum IgA**

Comparison between percentage of multimeric vs monomeric total IgA as estimated using HP-SEC or ELISA for same panel of 24 sera as in Figure 4E. Cutoff for calculating % polymeric IgA in SEC experiments was estimated on comparison with IgA-J measurements as shown in Figure S5B.
